# *Cis*-regulatory evolution shapes dehydration response in a desert-adapted house mouse

**DOI:** 10.64898/2026.03.13.710177

**Authors:** Miles Whedbee, Katya L. Mack

## Abstract

Deserts are among the most extreme environments on Earth. High temperatures and a lack of water impose powerful selective pressures on desert species, offering an opportunity to investigate the genetic basis of local adaptation. Despite the unique challenges of desert living, house mice (*Mus musculus domesticus*), a species native to Western Europe, have recently colonized the Sonoran Desert in North America within the last 400-600 generations. House mice from the Sonoran Desert show phenotypic differences consistent with adaptation to water scarcity, including maintaining weight better under water stress than non-desert mice. To investigate the genetic basis of the physiological responses to water deprivation, we compared gene expression responses of desert house mice and an interfertile non-desert dwelling subspecies (*M. m. musculus*) and their F1 hybrids after 72 hours without water access. First, we show that desert and non-desert mice exhibit highly divergent transcriptional responses to water deprivation across three tissues (hypothalamus, liver, and kidney). Then, by surveying allele-specific expression in intersubspecific hybrids between desert and non-desert mice, we uncover *cis-*regulatory differences driving changes in the transcriptional response to dehydration (e.g., *cis*-by-environment interactions). These *cis-*regulatory changes were highly tissue-specific, consistent with modular regulatory changes shaping expression divergence. Intriguingly, we find that genes with *cis-*regulatory differences induced by water access were involved in the arachidonic acid pathway, a primary adaptation pathway across many desert species, and lipid metabolism. Finally, our results highlight several candidate genes of interest for understanding rapid adaptation to desert living. Together, our results identify context-dependent *cis*-regulatory evolution as a key contributor to variation in dehydration response and a potential mechanism facilitating rapid adaptation to extreme environments.

## Introduction

A long-standing goal in evolutionary biology is understanding the genetic basis of adaptation. Desert environments represent an exciting opportunity to investigate the genetics of adaptation, because they impose extreme and well-defined selective pressures, including chronic water limitation, high temperatures, and intense solar radiation. Among the most significant of these stressors is acute and chronic dehydration. Many desert mammals have specialized physiological adaptations to dehydration, distinct from plastic responses observed under dehydration in non-desert species^1–3^. With the rising threats of desertification and drought associated with climate change^4,5^, understanding adaptations to arid environments is increasingly important.

Adaptations to limited water in desert environments are complex and wide-ranging, including specialized renal anatomy and physiology that minimizes water loss^6,7^, enhanced reliance on metabolic water to compensate for reduced environmental water availability^1,8^, and behavioral modifications^9^. While there has been great progress characterizing the physiological and morphological traits that enable desert mammals to survive in harsh arid environments, progress toward understanding the genetic basis of these desert phenotypes has been comparatively slower^3^. Recently, comparative genomic and population genomic approaches have begun to identify loci and pathways associated with desert adaptation, including signatures of positive selection at genes involved in kidney function, osmoregulation, and metabolism^10–16^. Still, comparative genomic approaches alone are often limited in their ability to establish mechanistic or causal links between genetic variation and adaptive phenotypes, particularly for complex, environmentally mediated traits such as dehydration tolerance. Transcriptomics can be a complementary approach for identifying genes and pathways associated with desert adaptation^12,17–20^. Experimental manipulations of water access in desert-adapted species have identified genes and pathways involved in governing transcriptional responses to dehydration^21–25^. Common garden experiments comparing transcriptional responses to water stress in closely related desert and non-desert species can identify divergent gene regulatory responses associated with dehydration tolerance in desert-adapted lineages^23^. However, comparative transcriptomic approaches, while powerful for identifying candidate genes and pathways for desert adaptation, can also be confounded by factors like differences in cell type abundances or developmental timing between individuals or species. Moreover, differential expression analyses alone cannot pinpoint whether divergent gene expression reflects direct regulatory changes at individual genes or upstream changes in gene regulation, making it difficult to identify the specific molecular changes underlying adaptive divergence.

House mice represent a powerful system with which to study recent climate adaptation, including to deserts^23,26^. Native to Western Europe, house mice colonized the Americas approximately 400-600 generations ago. In this time, house mice have invaded a wide range of environments in North and South America, where they show evidence of rapid environmental adaptation, including changes in behavior, body size, and physiology^27–33^. Among the most challenging of these environments is the Sonoran Desert, a hot and arid region of North America characterized by extreme heat and seasonally absent surface water^23,29^. Colonization of the Sonoran Desert likely imposed strong selective pressure on house mice, potentially driving local adaptation through changes to osmoregulation and water balance^23^. Supporting this idea, lab-born descendants of wild mice collected from the Sonoran Desert show phenotypic differences relative to non-desert populations suggestive of rapid adaptation to desert-living, including reduced water consumption, reduced weight loss under water stress, and differences in blood chemistry related to osmoregulatory function^23^. Moreover, comparisons of gene expression in the kidney between a desert and non-desert house mouse strain revealed widespread transcriptional differences following dehydration, supporting evolved differences in how desert mice respond to water stress^23^. Together, these findings suggest house mice in the Sonoran Desert have rapidly evolved to cope with periods of water scarcity, including modifications to their short-term response to dehydration.

We reason that adaptation to the Sonoran Desert in house mice may involve gene regulatory changes that alter the plastic response to dehydration. To examine this, here we investigate gene expression differences between lab-born descendants of desert-dwelling *Mus musculus domesticus* from Tucson, Arizona and a non-desert dwelling strain of the closely related subspecies *M. m. musculus* across three tissue types: kidney, liver, and hypothalamus. First, we compare gene expression in desert and non-desert mice under two hydration treatments: (1) free water access, and (2) water deprivation. Then, we generate F1 hybrids between desert and non-desert strains and quantify allele-specific expression (ASE) to characterize gene regulatory divergence underlying expression differences. ASE is a powerful approach that enables us to distinguish *cis-* (local) from *trans-* (distant) regulatory changes: in F1 hybrids, both alleles are exposed to a shared *trans* environment, such that allele-specific differences reflect *cis-*regulatory divergence, whereas expression differences observed between parental strains but not within hybrids are attributable to *trans* effects^34–37^. Importantly, this circumvents issues with differences in cell type abundances or developmental timing between species, as differences between the two alleles reflect *cis-*regulatory divergence acting within a shared cellular and environmental context. Finally, we compare ASE across water treatments to identify *cis-*regulatory changes underlying divergent responses to dehydration in desert mice. This comparative approach, known as differential allele-specific expression (diffASE), can be powerful for connecting context-specific traits or behaviors and *cis*-regulatory variation^38–41^.

Altogether, our results suggest an important role for *cis-*regulatory evolution in mediating evolved responses to water scarcity in desert house mice. We identify candidate genes and pathways also identified in long-established desert specialists, suggesting potentially convergent mechanisms underlie desert adaptation across diverse systems.

## Results and Discussion

### Mice from the Sonoran Desert lose less body weight under water stress

Decreased water availability can act as a strong selective pressure for plastic responses that ameliorate water stress. We reasoned that desert-dwelling house mice may have evolved divergent gene regulatory responses as a mechanism to cope with short periods of dehydration. We began by exploring the physiological response of a mouse strain derived from the Sonoran Desert (Tucson, Arizona: TUCC) and that of a control strain from a non-desert locality (Prague, Czech Republic: PWK), and their F1 hybrids to water stress (PWKxTUCC)(see Methods; Figure 1A). For our experimental group, we removed water entirely for a period of 72 hours from mice of all three genotypes (TUCC, PWK, PWKxTUCC). A second control group was provided unlimited water (water *ad libitum*). Consistent with previous results^42^, the desert-derived strain lost significantly less body weight after 72 hours of water stress compared to the non-desert strain (Mann-Whitney U, *p*=0.0078; median proportion weight maintained: TUCC: 0.78, PWK: 0.75; Figure 1B). Hybrids between PWK and TUCC lost a higher proportion of their body weight than TUCC and a similar amount to PWK (PWKxTUCC median proportion weight maintained: 0.76, TUCC vs. PWKxTUCC, Mann-Whitney U, *p*=0.0078; PWK vs PWKxTUCC, *p*=0.48; Figure S1). Weight loss is a marker of condition in animal husbandry, and monitoring weight loss is a classic approach for comparing the effects of dehydration^1,43–45^. Reduced weight loss in Sonoran desert mice is suggestive of enhanced tolerance for water deprivation or buffering against short periods of water stress.

**Figure 1.**
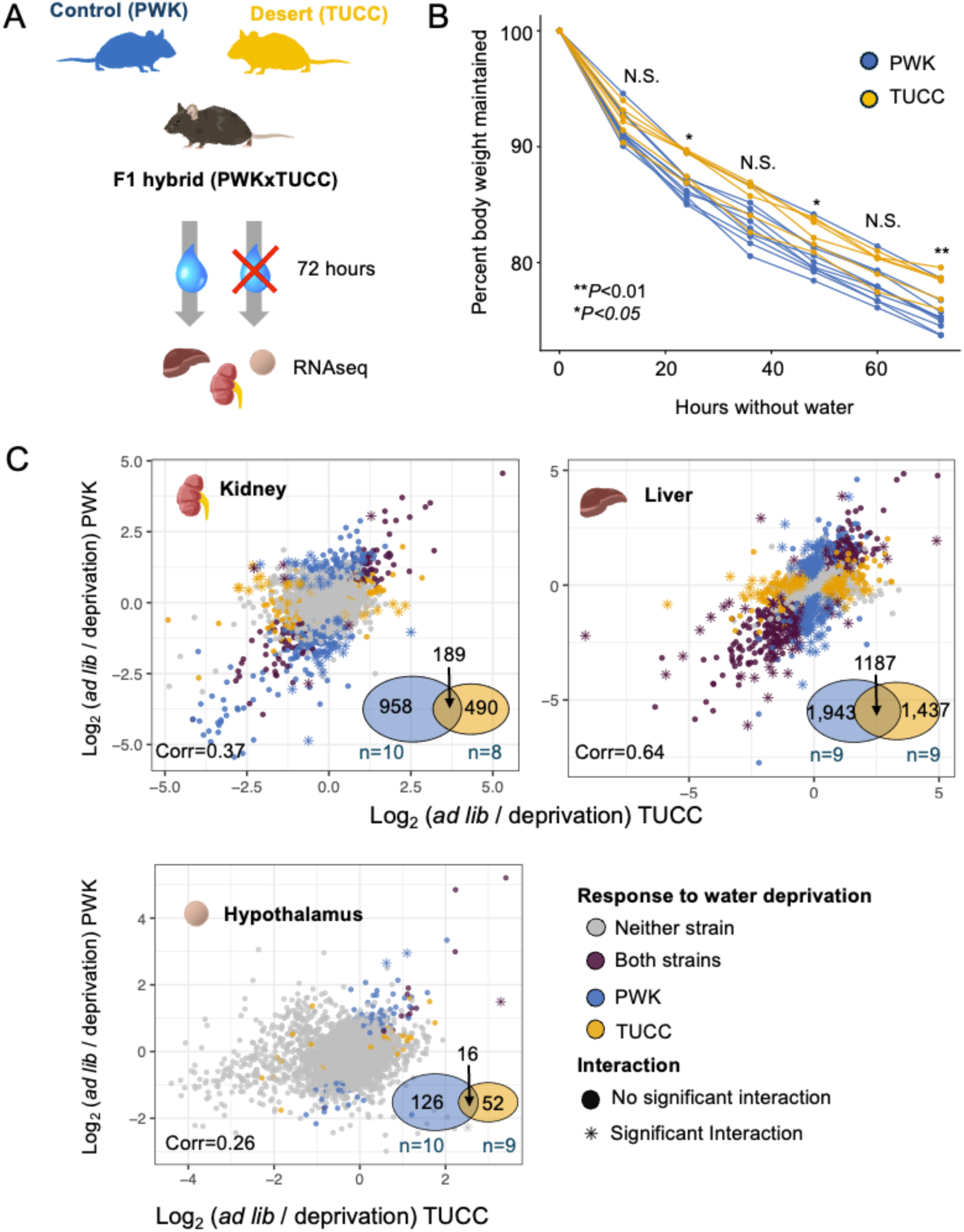
Highly divergent responses to water access between a desert and non-desert mouse strain. **A.** Mice from a desert (TUCC) and non-desert (PWK) strain and F1 hybrids were assigned to treatments groups, either 72 hours of water deprivation or 72 hours of unlimited water. After 72 hours, tissue was collected from each individual for RNAseq. **B.** TUCC mice maintained significantly more weight after 72 hours of water restriction (Mann-Whitney U tests). **C.** The transcriptional response to water deprivation in PWK vs. TUCC. Scatterplots compare log_2_ fold changes between water treatments across genotypes. Each point represents a gene. Genes with differential expression between water treatments are highlighted (blue = significant in PWK, gold = TUCC, both strains = purple). Significant interactions between genotype and treatment are designated by stars. Venn Diagrams show the number of genes that were responsive to water deprivation for each tissue type in PWK and TUCC.

### Expression divergence between desert-dwelling and non-desert dwelling mice

To measure the response to water stress on the transcriptional level, we generated RNAseq data from each genotype-treatment combination for 3 tissues selected for their role in water homeostasis: hypothalamus, liver, and kidney (Figure 1A). Whole-body homeostasis is maintained in large part by transport processes in the kidney, and the hypothalamus detects blood osmolarity and is involved in regulating thirst via the posterior pituitary gland^46^.

First, we compared the gene expression differences between the desert-derived TUCC strain and non-desert PWK strain. Across tissues, we found that 11,747 genes were differentially expressed between the TUCC and PWK mice with free water access and 11,089 genes were differentially expressed between TUCC and PWK after water deprivation (54% and 51% of genes surveyed)(DESeq2, Wald test, FDR<0.05)(Table S1). The majority of these expression differences were tissue-specific (65% and 68% in the *ad libitum* and water deprivation treatment, respectively). Many of the differentially expressed genes between the TUCC and PWK strains were also unique to a treatment group (42% of significant genes). Overall, gene expression differences between these strains were found to be highly tissue- and context-dependent.

### Divergent transcriptional responses to water deprivation in desert-dwelling and non-desert dwelling mice

Given the differing physiological response to water stress between the two mouse strains shown through differential weight loss, we examined the transcriptional response to water deprivation in each strain by comparing expression across water treatment groups (i.e., 72 hours of water deprivation vs. water *ad libitum*). We identified 4,057 and 3,160 genes with a significant transcriptional response to water deprivation in PWK and TUCC, respectively (FDR<0.05), with a greater proportion of genes differentially expressed in the non-desert lineage, consistent with a stronger transcriptional response to water stress. Regardless of strain, the fewest genes showed significant responses to treatment in the hypothalamus (194 genes total) and the most in the liver (4,567 genes total)(Figure 1C). A similar number of genes were upregulated vs. downregulated in response to water deprivation in most comparisons, with the exception of the TUCC hypothalamus, where the majority were upregulated after water deprivation (72%, 50 vs. 19 genes). Relatively few genes were differentially expressed in more than one tissue (205 in TUCC, 348 in PWK).

Comparing the transcriptional response to water deprivation between strains identified largely unique gene sets (Figure 1C), suggesting differences in the evolved dehydration response.

While these strains showed largely divergent responses to dehydration, 1,350 genes were identified in both, representing the shared transcriptional response to dehydration between these two strains. Many of these genes have previously described roles in water balance. For example, the gene *Stc1* was strongly upregulated in the kidney in both strains in response to water deprivation. *Stc1* is a gene involved in metabolism and dehydration response, coordinated by antidiuretic hormone (ADH)^15,47^. *Ptgs2*, also upregulated in the kidney in both genotypes, is induced during dehydration in the renal medulla, and disruptions to this gene impair urine concentrating ability^48,49^.

Testing specifically for interactions between strain and water treatment, we identified 389 genes with significant evidence for genotype-by-environment interactions across tissues (Figure 1C). Genotype-by-environment effects were tissue-specific, with no genes showing significant interactions in more than one tissue type. We found that significant gene-by-environment interactions were enriched for gene ontology (GO) terms including terms related to regulation of cell cycle (e.g., kidney: regulation of cell cycle process, *q*=2.94x10^-4^; mitotic cell cycle process, *q*=2.66x10^-3^; cell cycle phase transition, *q*= 7.29x10^-3^) and stress (e.g., response to stress, *q*=6.75x10^-2^).

### Evidence for gene regulatory changes in *cis* and *trans* between desert and non-desert mice

To investigate gene regulatory differences between desert and non-desert dwelling mice, we created F1 hybrids between TUCC and PWK. In F1 hybrids, alleles from both parents are exposed to the same *trans* environment and differences in expression between alleles in a hybrid (i.e., allele-specific expression) can be attributed to *cis*-regulatory differences between species^34,50^ (Figure 2A). Consequently, comparisons of parental expression ratios and that of allele-specific expression in hybrids can be used to infer *trans* divergence^34^. By comparing F1 hybrids across water treatments, we can also examine the impact of water restriction on gene regulatory differences between desert and non-desert mice.

**Figure 2.**
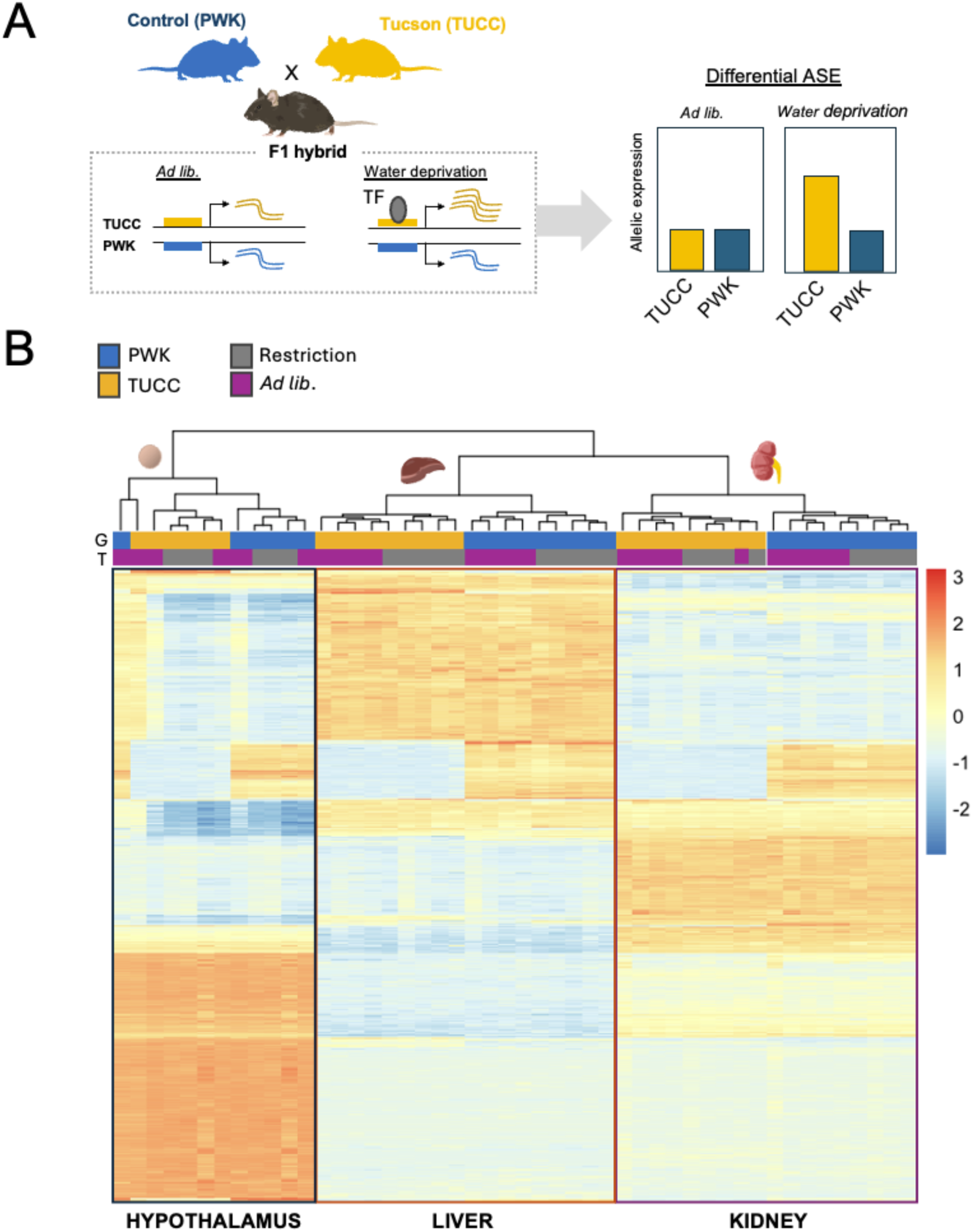
Allele-specific expression across three tissue types reveals *cis-*regulatory divergence. **A.** A schematic showing allele-specific expression (ASE) and differential allele-specific expression (diffASE). Allele-specific expression in F1 hybrids between PWK and TUCC is evidence of *cis-* regulatory divergence between strains. In this example, allele-specific expression is observed only after water deprivation, due to the binding of a context-specific transcription factor. **B**. A heatmap of the most variable genes (1,000) following variance stabilizing transformation of allele-specific read counts. Hierarchical clustering groups by tissue (hypothalamus [left], liver [middle], kidney[right]) and genotype (G: TUCC [gold], PWK [blue]), and incompletely by treatment (T: water deprivation [grey], water *ad libitum* [purple]).

We obtained an average of 32,135,867 allele-specific reads per sample, allowing us to test 17,002 genes for allele-specific expression across tissues and treatments (between 12,007-15,281 per tissue-treatment). Allele-specific expression values clustered samples strongly by tissue type (Figure 2B). Within a tissue, allele-specific libraries clustered largely by genotype, and incompletely by water treatment (Figure 2B). We detected between 2,725-6,401 genes with evidence for allele-specific expression (“ASE genes”) per tissue, indicative of *cis-*regulatory differences at these genes (DESeq2, FDR<0.05). The greatest number of ASE genes were detected in the kidney (Table S2, S3). The majority of ASE genes were tissue-specific (∼60%), but a subset were shared across all tissues (10% and 11% of ASE genes in the control and water deprivation treatment, respectively).

Comparisons of the distribution of F1 hybrid allelic ratios (log_2_[PWK allele / TUCC allele]) to the ratios of parental expression (log_2_[PWK *ad libitum* / TUCC *ad libitum*] expression) indicated the presence of both *cis* and *trans* differences contributing to expression divergence between species (Figure S2, Spearman’s *rho*=0.45-0.67). Comparisons in the water deprivation treatment were similar (Spearman’s *rho*=0.46-0.72). Both *cis-* and *trans-*acting gene regulation can be sensitive to environmental variation^37,51^, thereby affecting plastic transcriptional responses that allow organisms to adjust to changing conditions. Comparing between treatments, we observed significant shifts in magnitude for both *cis* and *trans* effects, indicating dehydration affects expression divergence through both *cis-* and *trans-* acting mechanisms (Figure S3, Wilcoxon test, *P*-values all < 2.2 x 10^-16^). We then asked whether *cis* or *trans* differences between TUCC and PWK were affected more by dehydration, again by comparing differences in log_2_ fold changes as proxies for effect size. In all tissues, we found that *trans* differences were greater across water treatments than *cis* changes (Figure S3, Wilcoxon rank sum tests, *P*-values all < 1.45 x 10^-10^). This is consistent with other studies which have also indicated a large role for *trans-* regulatory differences in plastic responses^31,51–55^.

### Extensive treatment-specific *cis*-regulatory changes in desert mice associated with dehydration response

*Cis-*regulatory changes play a predominant role in adaptation via changes in gene regulation^56–60^. As *cis*-regulatory mutations are often predicted to be less pleiotropic than protein-coding changes^37,56,61^, *cis*-regulatory variation may provide an important substrate for adaptation through context-specific modulation of gene expression in response to environmental stress. By comparing allele-specific expression across water treatments (i.e., water *ad libitum* vs. deprivation), we can identify *cis-*regulatory differences at specific genes between desert and non-desert mice in response to dehydration (*cis*-by-environment interactions) (Figure 2A).

Further, we reason that *cis-*regulatory differences revealed or modified differently under water stress in desert mice may play an important role in driving differences in physiological responses to water scarcity in desert mice.

To examine the extent to which *cis*-regulatory effects were shared or treatment-dependent, we compared ASE genes identified after water deprivation versus under water *ad libitum*. Between 53%-67% of ASE genes were shared across water treatments per tissue. ASE values were also correlated across treatments (Spearman’s rho=0.56-0.63, all *p-*values < 2.2 x10^-16^). Altogether, this suggests that while many *cis-*regulatory changes are robust to water treatment, a substantial fraction show context-specific effects. Given this observation, we sought to formally identify *cis*-by-treatment interactions by testing for differences in allelic expression ratios between the mice under water *ad libitum* vs. water deprivation (e.g., PWK/TUCC_deprivation_ vs. PWK/TUCC*_ad libitum_*) (Figure 2A; see Methods). These are genes in which allelic imbalance differs significantly between hydration conditions, indicating the presence of *cis-*regulatory changes driving plastic responses. Across all tissues, 483 genes were identified as having differential allele-specific expression (“DiffASE”) between treatments (DESeq2, FDR<0.05)(Figure 3A, full list in Supplemental File S1). The majority of these genes showed a difference in magnitude of allelic balance rather than a change in direction of bias across treatments (Kidney 190/241, Liver: 177/262, Hypothalamus: 5/6).

**Figure 3.**
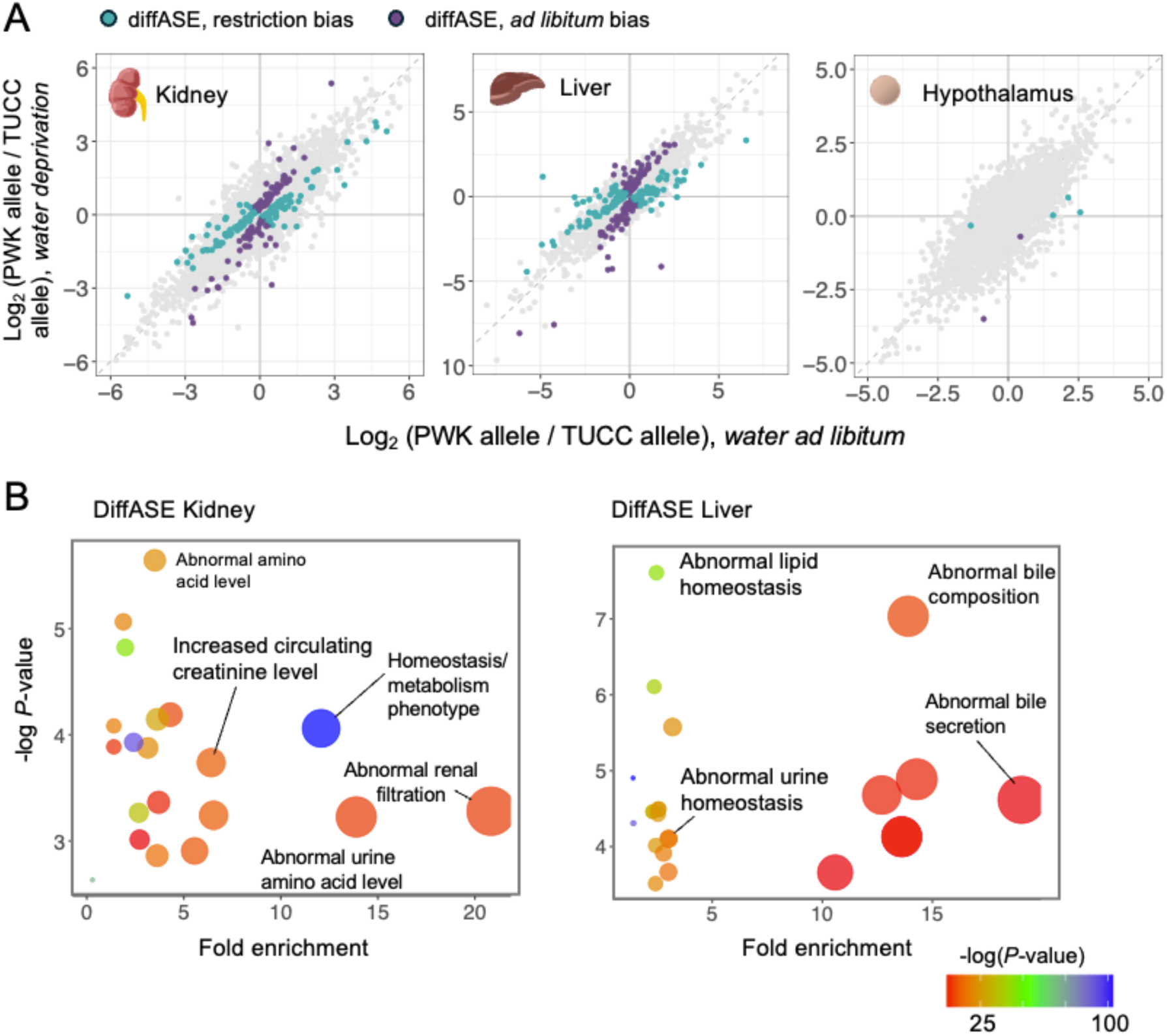
Context-specific *cis-*regulatory differences between desert and non-desert mice under differential water access. **A.** Scatterplots comparing the distribution of allelic ratios between water treatment conditions. Each point represents an individual gene. Highlighted are diffASE genes (purple points are genes with a larger difference between TUCC and PWK alleles after water restriction; blue points are genes with a larger difference between TUCC and PWK alleles for water libitum). **B**. Phenotype enrichments for diffASE genes in the kidney (left panel) and liver (right panel).

Only 26 genes showed diffASE in more than one tissue, meaning that these treatment-specific *cis-*regulatory differences were generally tissue-specific. Notably, we identified very few diffASE genes in the hypothalamus (6 genes: *Mustn1, Hsd3b2, Serpina1b, Slc27a5, Acaa1b, Hpn*).

Parental expression comparisons also indicated a more limited response to water treatment and few genotype-by-treatment effects for hypothalamus compared to other tissues (see Figure 1C). These results are consistent with more limited divergence between desert and non-desert mice in the strain-specific water stress response for hypothalamus compared to liver and kidney.

### Water treatment-specific *cis*-regulatory differences are associated with arachidonic acid and fat metabolism

While ASE genes will include many genes with strain-specific divergence unrelated to desert adaptation or water physiology, we reasoned that diffASE across treatments should be enriched for genes involved in dehydration response. To investigate this, we looked for evidence of functional enrichment in the diffASE gene set. We found that diffASE genes in the kidney were strongly enriched for genes with mutant phenotypes related to renal physiology (e.g., abnormal urine homeostasis, abnormal urine uric acid level, abnormal renal filtration, crystalluria) and homeostasis (e.g., abnormal amino acid level, increased circulating creatinine level, uremia)(full list in Supplemental Files S1). In the liver, diffASE genes were also enriched for phenotype terms related to the urinary system and urine homeostasis (e.g., abnormal renal/urinary system physiology, abnormal urine homeostasis), in addition to metabolism and homeostasis phenotypes (e.g., abnormal lipid homeostasis, abnormal lipid level, abnormal cholesterol homeostasis, abnormal bile composition) (Figure 3B). Adaptation to water stress in desert mammals often involves changes in renal phenotypes like urine homeostasis and water absorption (e.g., amount of urine produced, water reabsorption from kidney, changes in filtration rates) and metabolic phenotypes like fat metabolism^3,7^.

Examining pathway enrichments, we found that diffASE genes were enriched for several pathways, including the arachidonic acid metabolism pathway (Reactome pathways, R-MMU-2142753; kidney: FDR=1.18 x 10^-2^, liver: FDR=1.60 x 10^-2^), cytochrome P450 (R-MMU-211897; kidney: FDR=1.10 x 10^-2^, liver: FDR=7.64 x 10^-4^), fatty acid metabolism (R-MMU-8978868; kidney: FDR=1.04 x 10^-4^, liver: FDR=7.41 x 10^-7^), metabolism of lipids (R-MMU-556833 kidney: FDR=1.27 x 10^-4^, liver: FDR=1.89 x10^-8^), and cholesterol biosynthesis (R-MMU-191273; liver: FDR=9.62 x 10^-4^)(Supplemental Files S1). Intriguingly, the arachidonic acid metabolism pathway has been found to be a primary adaptation pathway across a wide range of desert species^3,11,62–64^. Changes in this pathway in desert mammals may affect water-retention mechanisms^3^. Our analysis highlighted 11 genes in the Arachidonic Acid metabolism pathway with diffASE in the liver or kidney (liver: *Akr1c20, Cbr1, Cyp2j6, Gpx1, Cyp2u1, Cyp4a12a*; kidney: *Cyp4a14, Cbr1, Akr1c21, Cyp4b1, Dpep1, Cyp4a12a*) representing >7-fold enrichment in both tissues. For genes in this pathway, we see both instances of greater *cis-*regulatory divergence under water restriction as well as reduced divergence under water restriction between strains (Figure 4). The cholesterol biosynthesis pathway, which showed a >12-fold enrichment for the liver, has also emerged as a potential adaptive target in other desert systems^22,65,66^. Genes in this pathway with diffASE have also been identified as potential candidate genes for desert adaptation in previous studies (Table S4).

**Figure 4.**
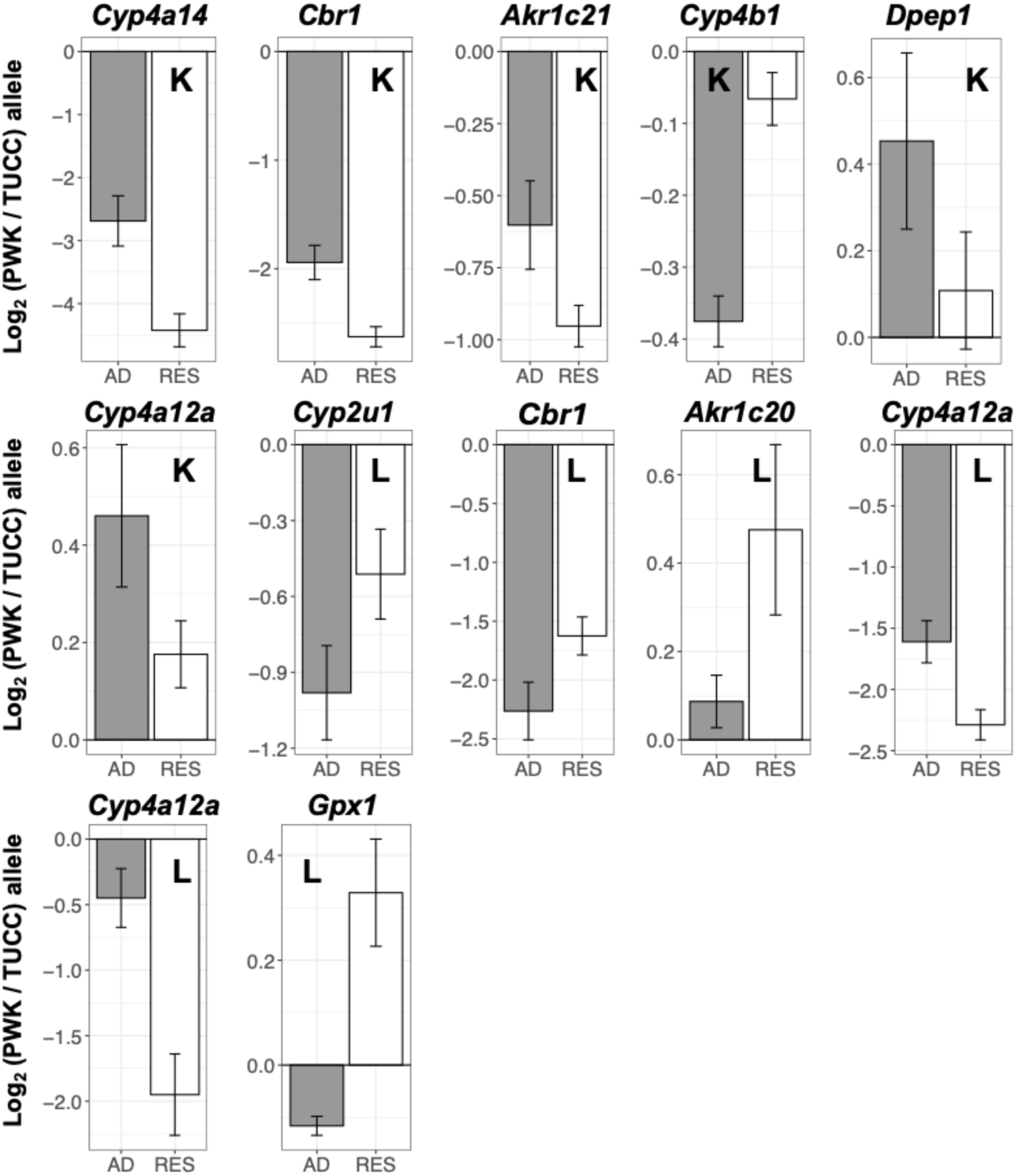
Differential allele-specific in the arachidonic acid metabolism pathway. Genes in the arachidonic acid metabolism pathway (R-MMU-2142753) with evidence of diffASE between the desert TUCC strain and the non-desert PWK strain. Bar plots show average log_2_ fold changes between alleles for *ad libitum* water (“AD”, grey bars) and water restriction (“RES”, transparent bars). Expression in the kidney is designated with a “K” and liver with an “L”.

Looking specifically at the 26 genes with diffASE in more than one tissue, we found that this set of genes was enriched primarily for metabolic process terms (e.g., fatty acid metabolic process, FDR=1.19 x 10^-6^; lipid metabolic process, FDR=5.74 x 10^-5^; acetyl-CoA metabolic process, FDR=1.68 x 10^-2^). Genes involved in fat metabolism have also been highlighted as potential candidates for desert adaptation in several other desert systems^3,10,67–69^. Genes involved in fat metabolism have been suggested to contribute to dehydration tolerance via enhanced use of metabolic water from fat^3^.

### Evidence for allelic induction associated with dehydration response

Next, we examined genes with evidence of allelic induction or repression^39,40^. These differ from diffASE genes, in that we are interested in comparing the expression of an individual allele across treatments (e.g., induction or suppression of desert TUCC allele in water deprivation vs. water *ad libitum*), similar to a traditional differential expression analysis.

In the kidney, we observed 14 diffASE genes with evidence of TUCC allelic induction under water deprivation and 32 genes with TUCC allelic suppression. In the liver, 24 diffASE genes showed evidence of induction and 86 showed evidence of suppression of the TUCC allele. This analysis again highlighted genes related to arachidonic acid metabolism. Five members of the Cytochromes P450 gene family (*Cyp2j13, Cyp51, Cyp2c23, Cyp2d9, Cyp4a14*) showed significant induction or repression of the TUCC allele under water deprivation in the kidney and 4 in the liver (*Cyp2c23, Cyp2d10, Cyp2u1, Cyp4v2*). *Cyp4a14* showed >2 log_2_ fold change greater induction of the TUCC allele vs. the PWK allele under water deprivation in the kidney.

*Cyp4a14* is involved in 20-HETE synthesis, and mutants affect renal excretory capacity^70,71^. Investigating specifically genes where the TUCC allele was induced or repressed but the PWK allele did not show changes in expression across conditions (32 genes), we found enrichments for mutant phenotypes including renal/urinary system (MP:0010768, *q*=0.036), cholesterol homeostasis (MP:0005278, *q*=0.041) and bile composition (MP:0004773, *q*=0.012).

This analysis also identified cases of induction or repression of the PWK allele (65 and 41 diffASE genes in the liver and kidney, respectively). DiffASE genes where only the PWK allele changes expression across conditions may also be of interest, as these may include cases where plastic responses are buffered in desert mice to tolerate dehydration. For the kidney, these genes were enriched for mutant phenotypes affecting the renal glomerulus, the network of small blood vessels located in the nephron of the kidney that acts as the primary filtration unit (e.g., abnormal glomerular mesangium morphology, *q*=0.02; renal glomerular protein deposits, *q*=0.01, renal glomerular immunoglobulin deposits, *q*=0.003).

## Conclusions

A primary challenge for mammals living in desert environments is extreme water scarcity. Here, we examined the response of house mice that have recently colonized the Sonoran Desert to short-term dehydration. Mirroring what was found previously^23^, we observed attenuated responses to water stress in the desert TUCC strain, with these mice maintaining more body weight and showing reduced expression divergence after 72 hours of water deprivation. Weight is a key indicator of condition that has previously been used to compare the overall effects of dehydration^43,72^.

At the transcriptional level, desert and non-desert mice displayed highly divergent gene expression responses to dehydration in the kidney, liver, and hypothalamus. Although a core set of genes responded similarly in both strains—representing a conserved plastic response to water deprivation—the majority of dehydration-responsive genes were strain-specific indicating differences in the dehydration response between desert and non-desert mice. Genotype-by-environment interactions were also tissue-specific, suggesting an organ-specific nature of divergent responses to water stress.

Using allele-specific expression (ASE) in F1 hybrids, we examined the regulatory basis of expression differences between desert and non-desert mice. We showed that both *cis-* and *trans-*regulatory divergence contribute to expression differences between desert and non-desert mice, with *trans* effects dominating overall plastic responses to dehydration. This is consistent with a growing body of work indicating that environmentally induced transcriptional plasticity is often mediated by *trans-*acting factors^31,51,73^. *Trans* effects (e.g., transcription factors) can play an important role in plastic responses because they can act as dynamic regulators of gene regulatory networks in the face of environmental stimuli. However, we also identified hundreds of genes with water treatment-specific *cis*-regulatory divergence, suggesting that environmentally induced *cis*-regulatory differences may contribute to adaptive responses. While we cannot conclude that these *cis-*regulatory changes are directly involved in dehydration tolerance or desert adaptation from these data alone, the enrichment of these genes for relevant phenotypes and pathways make them promising candidates.

Genes exhibiting water treatment-dependent *cis-*regulatory divergence were enriched for pathways involved in renal physiology, lipid metabolism, and notably, arachidonic acid metabolism. In mammals, arachidonic acid metabolites play key roles in renal blood flow, sodium transport, and urine concentration^74^. Regulatory changes affecting enzymes in this pathway, including cytochrome P450 genes (e.g., *Cyp4a14*^70^) may influence water retention and excretory capacity. Strikingly, the arachidonic acid pathway has repeatedly emerged as a target of selection in diverse desert lineages^3^ – including camels^62^, sheep^75^, deer^63^, and other rodents^11,13^ – suggesting that this pathway may have been repeatedly targeted during desert adaptation.

DiffASE genes were also enriched for lipid and fatty acid metabolic processes, including cholesterol biosynthesis. Lipid metabolism has been repeatedly implicated in desert adaptation across mammals^3^, potentially reflecting the importance of metabolic water production from fat oxidation during periods of water scarcity. In camels, alteration of the cholesterol biosynthesis pathway has been suggested as a mechanism to facilitate water retention in the kidney^22^.

Cholesterol biosynthesis genes also show differential expression in response to water regime across diverse desert lineages, supporting a role for cholesterol during dehydration^65^. The enrichment of lipid-related pathways among genes with water treatment-specific *cis-*regulatory divergence suggests that recent adaptation in Sonoran Desert house mice may parallel broader evolutionary patterns observed in long-established desert specialists. Characterizing how lipid levels change in this system under different water regimes may lend further insight into the role of these pathways in dehydration tolerance in house mice.

Overall, our results support a model in which rapid adaptation may involve context-dependent regulatory changes that reshape plastic transcriptional responses. Further, by having identified pathways involved in adaptation in desert specialists — particularly lipid and arachidonic acid metabolism — our results suggest a connection between recent, population-level adaptation in house mice and deeply conserved evolutionary solutions to life in arid environments.

## Methods

### Mouse husbandry and tissue collection

All experiments with animals were conducted with approval from the University of Kansas Institutional Animal Care and Use Committee. To examine responses to water deprivation in house mice after their invasion of the Sonoran Desert, we used a newly derived inbred line from mice captured in Tucson, Arizona (TUCC)^23,29^. Descended from wild-caught progenitors, this strain was generated through brother-sister mating at UC Berkeley. This strain was selected because it had been previously used in Bittner et al. 2021 to study physiological and transcriptional responses to water deprivation. Another strain from Tucson, Arizona is now also commercially available^33^. To maximize genetic distance for measurements for ascertaining allele-specific expression, TUCC male mice were crossed to a wild-derived strain of *Mus musculus domesticus*, PWK/PhJ (The Jackson Laboratory, strain #:003715, “PWK”) from a non-desert region as a non-desert control strain. PWK is a wild-derived inbred strain descended from wild mice trapped near Prague in the Czech Republic.

Adult male mice were randomly assigned to either the treatment (72 hours of water deprivation) or control group (water *ad libitum*). During the 72-hour experimental period, mice were housed singly and provided food *ad libitum*. Mice undergoing a period of water deprivation were weighed every 12 hours (Figure S1) and monitored for declining health. We observed results similar to what was observed previously in Bittner et al. 2021, where TUCC mice were found to lose less weight under water deprivation compared to another non-desert strain of *M. m. domesticus* from Edmonton, Canada. However, all mice lost a higher percentage of body weight on average in this study (Table S5). After 72 hours, mice were exposed to isoflurane until non-responsive and then euthanized via cervical dislocation. Left kidney, liver, and hypothalamus were immediately collected into RNAlater for RNAseq.

### RNA Extraction, Library Preparation, and Sequencing

RNA was extracted from three tissues—left kidney, liver, and hypothalamus—using TRIzol reagent (Invitrogen, 15596018) according to the manufacturer’s protocol. To avoid potential genomic contamination, RNA samples were treated with DNase I (Thermo Scientific, EN052) following the manufacturer’s protocol. Libraries were prepared using the NEB Ultra II Directional prep kit with the NEBNext Poly(A) Magnetic Isolation Module (New England Biolabs, Ipswich, MA, USA), pooled, and then sequenced on the Novaseq platform (2x150) through Admera Health. Samples information and number of reads obtained per sample are available in File S1. Due to issues with sample collection and library preparation for RNAseq, an unequal number of libraries were obtained for each genotype/tissue type (3-5 libraries genotype/treatment)(Table S6, File S1).

### Read mapping and parental gene expression analysis

Reads were trimmed for adapter sequences using the Cutadapt wrapper TrimGalore^76^. RNAseq reads were mapped with STAR (filter: --outFilterMultimapNmax 1)^77^ to the house mouse reference genome (GRCm39) downloaded from Ensembl. HTSeq-count was used to count reads overlapping exonic regions to produce gene-level count data^78^ based on gene annotations downloaded from Ensembl. Count data was then analyzed with DESeq2^79^ to identify differences in gene expression associated with (1) dehydration response (e.g., PWK water deprivation vs. PWK water *ad libitum*), genotype (e.g., TUCC water deprivation vs. PWK water deprivation), and the interaction between response and genotype (genotype * water treatment). Each tissue type was analyzed separately. *P-*values were corrected for multiple testing via the Benjamini and Hochberg method. We considered genes with an adjusted *P-*value (FDR) < 0.05 to be differentially expressed between comparisons. To examine transcriptome-wide patterns of gene expression for each tissue, we transformed expression values using variance stabilizing transformation and examined patterns via principal components analysis (PCA). PCA separated individuals by genotype in each tissue (Figure S4). We also observed separation by treatment in the liver on PC2 and separation based on treatment in the kidney (Figure S4).

### Whole-genome re-sequencing of TUCC and variant calling

DNA was extracted using the DNeasy blood and tissue kit (Qiagen, 69504) from TUCC tissue. Library preparation and sequencing was performed by Admera Health, using the Kappa HyperPrep kit, and sequenced to an average autosomal depth of 37.5x.

Genomic reads were downloaded from the Wellcome Trust Mouse Genome Project for PWK (https://www.sanger.ac.uk/data/mouse-genomes-project/)^80,81^. Reads from PWK and TUCC were mapped using BWA^82^ to the mouse reference genome. BCFtools^83,84^ was used for variant calling. We then filtered SNP calls for depth and quality (BCFtools filter: QUAL>20 && DP>10) and retained only homozygous sites. This list of variant positions was used as initial input for mapping with STAR (see below) and subsequently filtered further based on the presence of informative reads from RNAseq libraries, as described in “*Identifying allele-specific expression”* below. This resulted in a list of 2,663,253 informative sites for allele-specific expression analysis.

### Identifying allele-specific expression

To quantify ASE, we mapped reads from hybrid individuals to the house mouse reference genome with STAR. The WASP-implementation of STAR was used to reduce reference mapping bias, removing reads that do not map to the same location with the alternative allele^85,86^. We then quantified allelic reads per site using the GATK^87^ tool ASEReadCounter. To ensure a high-quality list of variants was being used for ASE quantification, we removed sites for which we did not obtain an overlapping TUCC and PWK read and then subsequently re-mapped our libraries using the same approach with these sites filtered out.

We then separated reads overlapping informative variants into allele-specific pools based on genotype (i.e., TUCC or PWK). HTSeq-count was used to count reads overlapping exonic regions, as described above. As ASE estimates can be less reliable at low read depths^88^, we filtered based on read counts, requiring a minimum of 20 reads per allele in a tissue. We then compared the reads mapped preferentially to either the TUCC or PWK allele using DESeq2, as described previously^41,51,89^ following a model incorporating allele (TUCC vs. PWK), treatment (water *ad libitum* vs. restricted), and sample information. As we had unequal sample sizes across treatments and genotypes, we randomly dropped samples to equalize power for comparisons within a tissue for ASE analyses. Previously published code is available on Github (github.com/malballinger/BallingerMack_PNAS_2023). We considered genes significant at an FDR threshold of 5%. The number of ASE reads obtained per sample can be found in Supplementary File S1.

### Enrichment analyses

Pathway and GO ontology enrichments were performed with GOrilla^90^ and Panther^91^ and phenotype enrichment analyses were performed with ModPhea^92^.

## Acknowledgment

We thank Michael Nachman for graciously sharing the wild-derived inbred strain used in this study. We thank Noelle Bittner and members of the Mack lab for feedback on the manuscript. We thank Joseph Ward for husbandry assistance.

## Funding

Research was supported by the National Institute of General Medical Sciences of the National Institutes of Health under Award Number R35GM154966 to K.L.M.

## Data Availability

All sequence data has been deposited online through NCBI Bioproject (PRJNA1432092). Additional supplemental material can be found in Supplemental File S1.

## Notes

### Competing Interest Statement

The authors have declared no competing interest.

