## Supplemental Tables and Figures for "*Cis*-regulatory evolution shapes dehydration response in a desert-adapted house mouse"

**Table S1.** Differentially expressed genes between TUCC and PWK.

| | Differentially expressed genes |
|---:|---:|
| Water deprivation, kidney | 5,055 |
| Water *ad lib.,* kidney | 6,653 |
| Water deprivation, hypothalamus | 4,508 |
| Water *ad lib*., hypothalamus | 4,061 |
| Water deprivation, liver | 5,990 |
| Water *adlib*., liver | 6,057 |

**Table S2.** Genes with ASE in each tissue.

| | Genes with ASE, water deprivation | Genes with ASE, water *ad lib.* | diffASE |
|---|---|---|---|
| Kidney | 6279 | 6401 | 241 |
| Hypothalamus | 2996 | 2725 | 6 |
| Liver | 4406 | 4425 | 262 |

**Table S3.** ASE across tissues, partitioned by effect size ($log_2$ fold change).

| Tissue and treatment | ASE (all) | ASE (>0.5) | ASE (>1) | ASE (>2) |
|---|---|---|---|---|
| Kidney, *ad lib* | 6401 | 3208 | 1539 | 485 |
| Kidney, water deprivation | 6279 | 3038 | 1474 | 459 |
| Liver, *ad lib* | 4406 | 2621 | 1303 | 407 |
| Liver, water deprivation | 4425 | 2550 | 1229 | 344 |
| Hypothalamus, *ad lib* | 2725 | 1527 | 742 | 241 |
| Hypothalamus, water deprivation | 2996 | 1689 | 752 | 224 |

**Table S4.** Cholesterol biosynthesis pathway [R-MMU-191273 ] genes with diffASE in the liver.

| Gene | Information from previous studies | Citation |
|---:|---:|---:|
| *Mvk* | Expression changes under different hydration conditions in the Arabian camel | Alvira-Iraizoz et al. 2021 |
| *Msmo1* | Positive selection in jerboas | Peng et al. 2023 |
| *Dhcr7* | Expression changes under different hydration conditions in the Arabian camel | Alvira-Iraizoz et al. 2021 |
| *Hmgcr* | Deletion in old world camel species | Guo et al. 2025 |
| *Mvd* | Expression changes under different hydration conditions in the Arabian camel | Alvira-Iraizoz et al. 2021 |
| *Acat2* | Expression changes under different hydration conditions in the Arabian camel | Alvira-Iraizoz et al. 2021 |

**Table S5.** Median weight lost after 72 without water in this experiment and Bittner et al. 2021.

| | Average percentage body weight maintained |
|---|---|
| TUCC (this study) | 78% |
| TUCC (Bittner et al. 2021) | 82% |
| PWK (this study) | 75% |
| Edmonton strain (Bittner et al. 2021) | 78% |

**Table S6. Number of libraries from each treatment/genotype.**

| Gentoype, treatment | Liver | Hypothalamus | Kidney |
|---:|---:|---:|---:|
| PWK, water restriction | 5 | 5 | 5 |
| TUCC, water restriction | 4 | 3 | 3 |
| Hybrid, water restriction | 4 | 3 | 4 |
| PWK, *ad lib.* | 4 | 5 | 5 |
| TUCC, *ad lib.* | 5 | 5 | 5 |
| Hybrid, *ad lib.* | 5 | 3 | 5 |

**Figure S1.** Body weight maintained under water restriction.

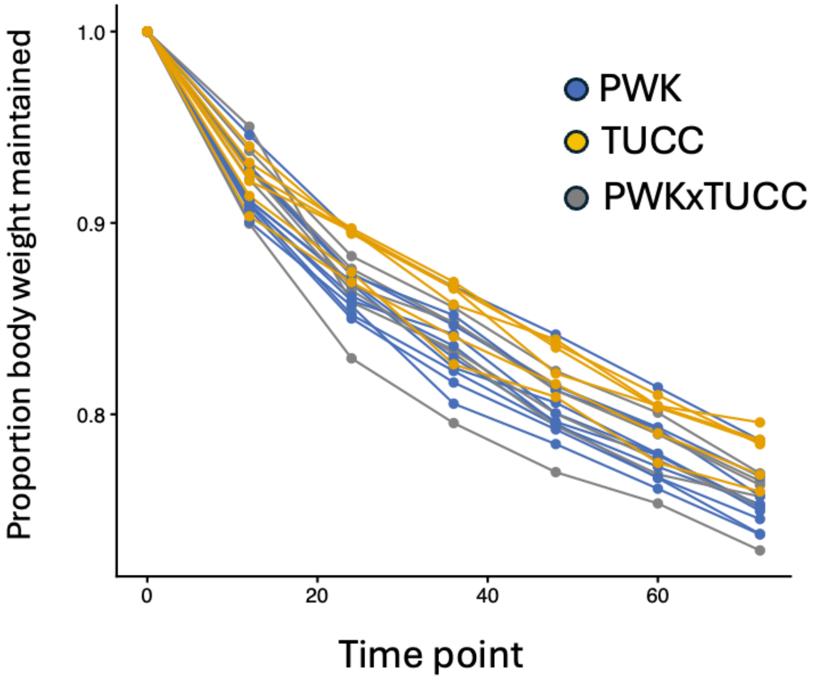

**Figure S2**. Scatterplot of the distribution of allelic expression in F1 hybrids and parental expression. Each point is a gene.

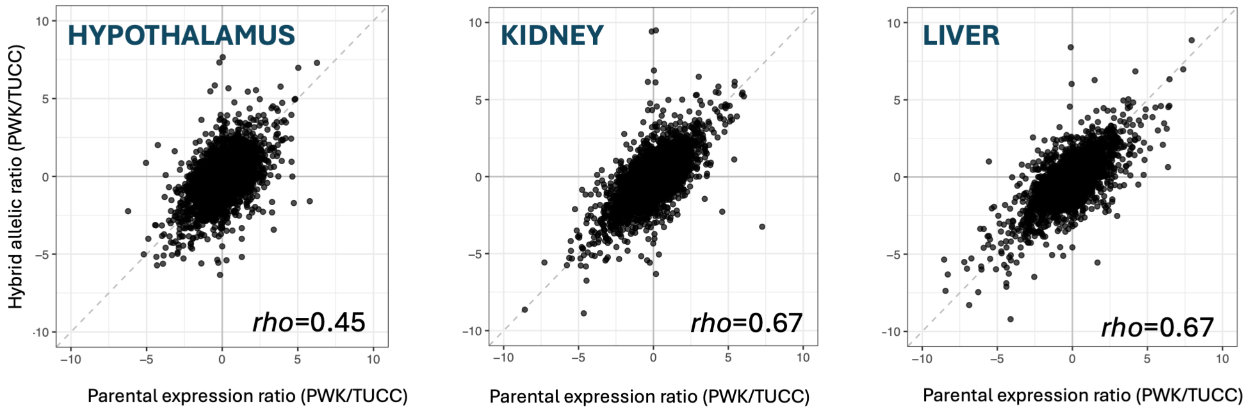

**Figure S3**. Distribution of effect size differences based on $log_2$ fold changes for *cis* (**A**) and *trans* (**B**).

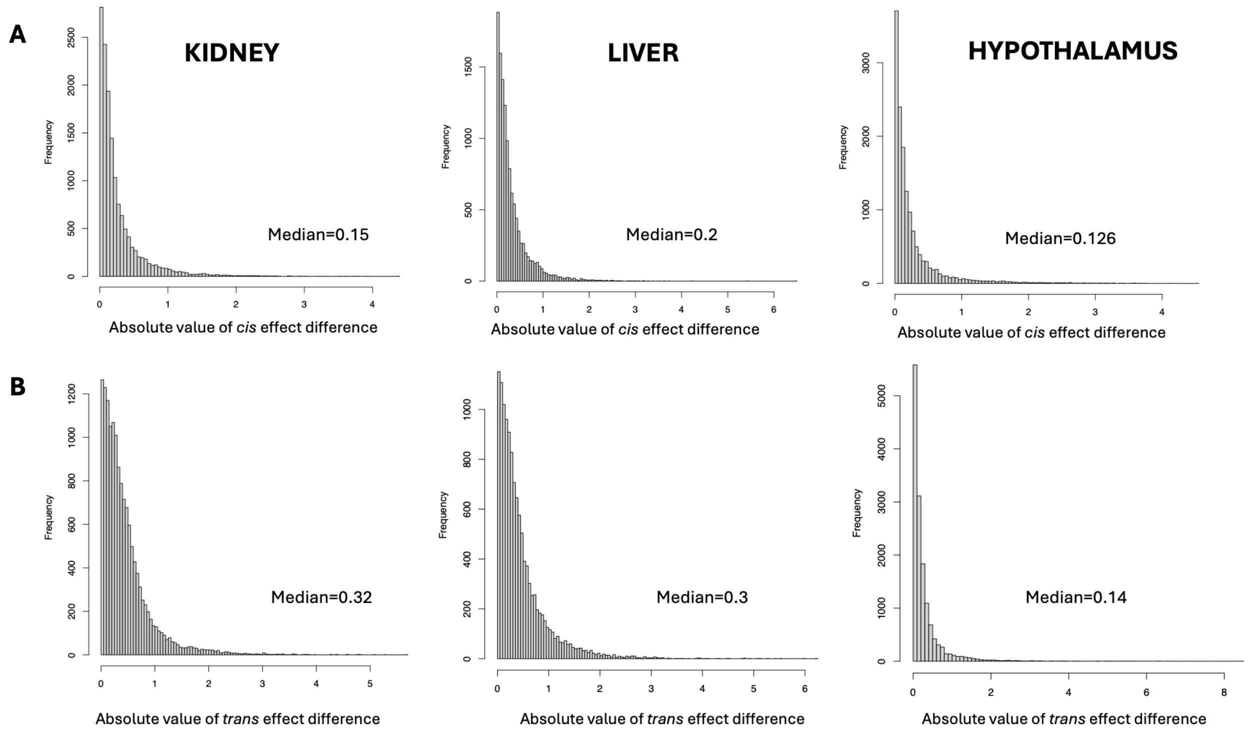

**Figure S4.** Principal component (PC) analysis of gene-wise mRNA abundance in TUCC and PWK (R = water restriction, A = water *ad libitum*)

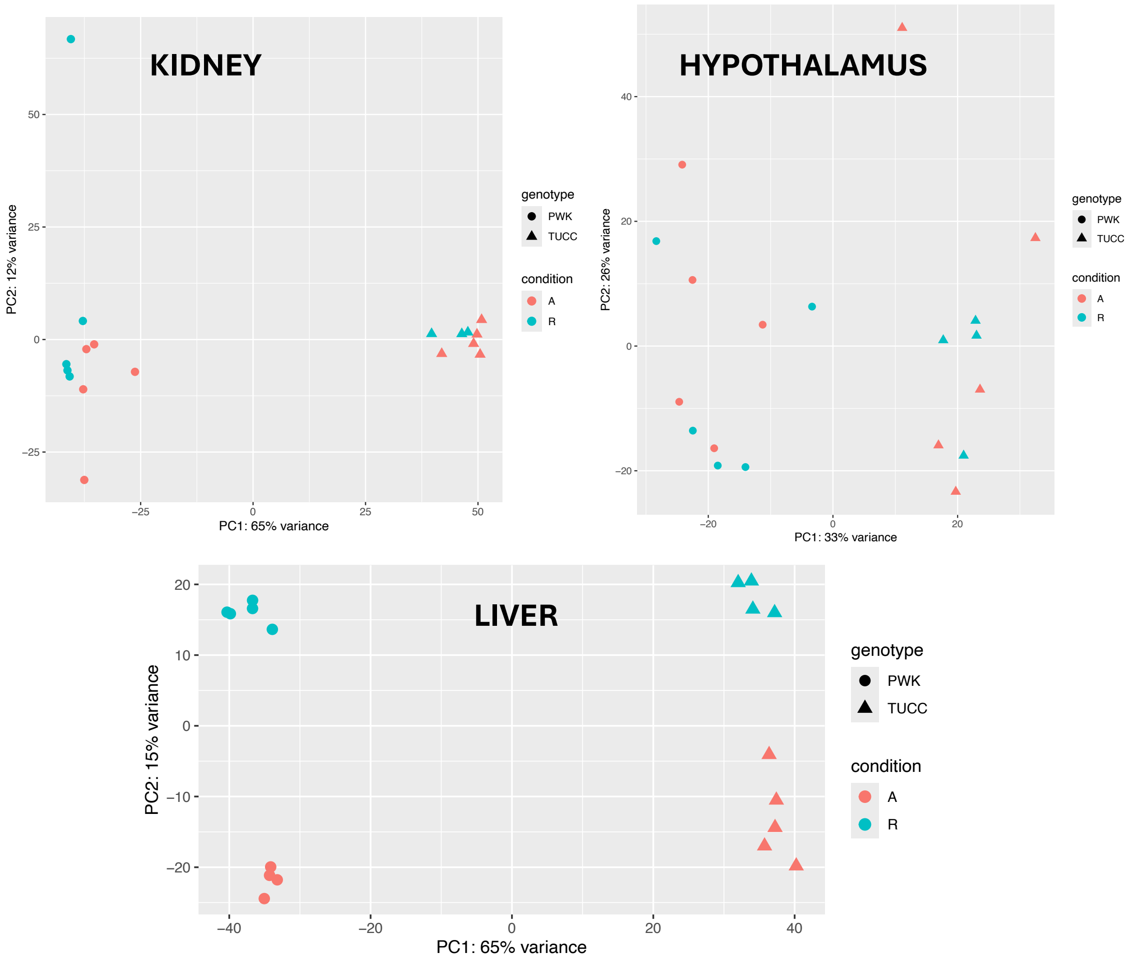
